# ABySS 2.0: Resource-Efficient Assembly of Large Genomes using a Bloom Filter

**DOI:** 10.1101/068338

**Authors:** Shaun D Jackman, Benjamin P Vandervalk, Hamid Mohamadi, Justin Chu, Sarah Yeo, S Austin Hammond, Golnaz Jahesh, Hamza Khan, Lauren Coombe, Rene L Warren, Inanc Birol

## Abstract

The assembly of DNA sequences *de novo* is fundamental to genomics research. It is the first of many steps towards elucidating and characterizing whole genomes. Downstream applications, including analysis of genomic variation between species, between or within individuals critically depends on robustly assembled sequences. In the span of a single decade, the sequence throughput of leading DNA sequencing instruments has increased drastically, and coupled with established and planned large-scale, personalized medicine initiatives to sequence genomes in the thousands and even millions, the development of efficient, scalable and accurate bioinformatics tools for producing high-quality reference draft genomes is timely.

With ABySS 1.0, we originally showed that assembling the human genome using short 50 bp sequencing reads was possible by aggregating the half terabyte of compute memory needed over several computers using a standardized message-passing system (MPI). We present here its re-design, which departs from MPI and instead implements algorithms that employ a Bloom filter, a probabilistic data structure, to represent a de Bruijn graph and reduce memory requirements.

We present assembly benchmarks of human Genome in a Bottle 250 bp Illumina paired-end and 6 kbp mate-pair libraries from a single individual, yielding a NG50 (NGA50) scaffold contiguity of 3.5 (3.0) Mbp using less than 35 GB of RAM, a modest memory requirement by today’s standard that is often available on a single computer. We also investigate the use of BioNano Genomics and 10x Genomics’ Chromium data to further improve the scaffold contiguity of this assembly to 42 (15) Mbp.

## Introduction

*De novo* genome assembly remains a challenging problem, especially for large and complex genomes. The problem refers to identifying partial and unambiguous overlaps between sequencing reads (which are orders of magnitude shorter than the target genome) to build longer, contiguous sequences (contigs) (Nagarajan and Pop 2013). If further linkage information is available, such as in the form of paired end reads or physical maps, these contigs may be ordered and oriented with respect to each other and reported as scaffolds, where there may be undetermined sequences (represented as ‘N’s) between contigs. It is practically accepted that assembly algorithms almost never reconstruct genomes in their full chromosomes (Paulino et al. 2015), and the quality of returned contigs and scaffolds are conventionally measured by the contiguity of the assembled sequences. Often assembly algorithms are also validated using data from resequencing experiments, where assembled sequences are compared against a reference genome for their correctness in addition to their contiguity (Gurevich et al. 2013).

Of particular interest in this study is resequencing data from human genome studies. The unbiased approach of *de novo* assembly of data from these experiments prior to comparison to a reference sequence is a valuable approach in detecting structural variants between individuals or between tumor and normal genomes (Li 2015; Mose et al. 2014). Even though it is substantially more computationally intensive to analyze sequencing data by assembling the reads first, gained specificity and the resulting savings in event verification efforts may justify the choice, but its heavy resource use also points to an area for improvement.

In this domain ABySS v1 was the first scalable *de novo* assembly tool that could assemble a human genome, using short reads from a high-throughput sequencing platform (Simpson et al. 2009). However, the feat required aggregating a large amount of memory distributed across a number of compute nodes communicating through a Message Passing Interface protocol. Although this enabling technology found applications in many large cancer cohort studies (Yip et al. 2011; Roberts et al. 2012; Pugh et al. 2013; Ley et al. 2013; Morin et al. 2013), its large memory footprint constituted a substantial bottleneck.

Of course this large memory footprint issue was not unique to ABySS v1, with many algorithms that can scale to assemble the human genome, including SOAP-denovo (Luo et al. 2012), SGA (Simpson and Durbin 2011), ALLPATHS_LG (Gnerre et al. 2010), MaSuRCA (Zimin et al. 2013) and DISCOVAR (Weisenfeld et al. 2014) all requiring around 1 TB of RAM, if not more, to accomplish this. To alleviate this bottleneck, Minia (Chikhi and Rizk 2013) and BCALM2 (Chikhi et al. 2016) algorithms introduce probabilistic data structures using Bloom filters (Bloom 1970) and minimizer hashing (Chikhi et al. 2014), respectively.

In ABySS v2, we follow the model of Minia, where sequence overlaps are inferred from a de Bruijn graph (Pevzner et al. 2001) representing an implicit Bloom filter.

As in ABySS v1, we catalogue all observed sequences of length k (k-mers, with k less than the read length), and store them in a Bloom filter. This representation of the k-mer spectrum of the input reads lends itself naturally to identify k-1 base pair overlaps, hence describe a de Bruijn graph.

Performance of sequence assembly algorithms is closely coupled with the sequencing technology used and the quality of the data they generate, with highly accurate long reads always being desirable. However, the genomics research landscape, especially cancer genomics studies, has been heavily dominated by the high throughput sequencing platforms from Illumina (San Diego, CA). Although longer (albeit noisier) sequences from Pacific Biosciences (Menlo Park, CA) instruments are proven to yield high quality *de novo* human genome assemblies (Chaisson et al. 2014; Pendleton et al. 2015), they come at a higher price relative to Illumina reads. The newer long read instruments from Oxford Nanopore Technologies (Oxford, UK) do not yet have the necessary throughput or data quality to be of utility in human genomics studies. As a result, most large cohort projects, as well as price-sensitive personalized medicine applications still use the Illumina platforms.

Another new technology, Chromium from 10X Genomics (Pleasanton, CA), generates sequencing libraries that localize sequence information on DNA fragments that are over 100 kb long. The technology employs microfluidics to isolate large fragments, which are sheared and barcoded separately, and prepared to sequencing libraries compatible with Illumina paired end sequencing. The barcodes can then be used to reconstruct the sequence of the long fragments from which they originate, providing valuable linkage information for assembly and scaffolding problems.

Further on the scaffolding problem, it was demonstrated in the original Human Genome Project (Lander et al. 2001) and other pioneering *de novo* sequencing projects that used Sanger sequencing data that linkage information from a physical map is very valuable in building highly contiguous assemblies. Although the approach is not favored in many studies, presumably for the additional cost that it brings, new optical mapping technologies, such as that from BioNano Genomics (San Diego, CA) represent intriguing opportunities.

In this paper we describe the details of the Bloom filter implementation in ABySS v2, and compare its performance with respect to the latest version of ABySS v1, as well as other scalable assembly pipelines, SOAPdenovo and SGA. We note that there are other algorithms that can build contigs from high throughput sequencing datasets, and we include comparison to DISCOVARdenovo, Minia and BCALM2 in that category, further contiguating their results using third party scaffolding algorithms, BESST (Sahlin et al. 2016), LINKS (Warren et al. 2015) and the scaffolding algorithm within the ABySS package. We further demonstrate how long range linkage information from Chromium reads and BioNano maps may improve scaffold contiguity of draft assemblies.

## Results

### ABySS 2.0.0 Assembly Algorithm

In ABySS 2.0.0, we have implemented a multi-stage *de novo* assembly pipeline consisting of unitig, contig, and scaffold stages. In the unitig stage, we perform the initial assembly of sequences according to the de Bruijn graph assembly paradigm (Pevzner et al. 2001). In the contig stage, we align the paired-end reads to the unitigs and use the pairing information to orient and merge overlapping unitigs. In the scaffold stage, we align the mate-pair reads to the contigs to orient and join them into scaffolds, inserting gaps of ‘N’ characters at gaps in coverage and unresolved repeats.

The main innovation of ABySS 2.0.0 is a Bloom filter-based implementation of the de Bruijn graph assembly algorithm that reduces the overall memory requirements of ABySS by an order of magnitude. While the original ABySS publication (Simpson et al. 2009) introduced a novel distributed approach for assembling large genomes on a cluster, the Bloom filter approach we describe here enables large genome assemblies to be run on a single machine with modest memory and achieves comparable results.

### Effect of Bloom Filter False Positive Rate

While a Bloom filter can be used to implement a highly compact representation of the de Bruijn graph, it comes with the caveat that it is a probabilistic data structure. In particular, a Bloom filter may return *false positives* when queried for the presence of particular *k*-mers in the graph. The false positive rate of a Bloom filter is determined by the size of the Bloom filter *m*, the number of elements inserted into the Bloom filter *n*, and the number of hash functions *h*, as first derived in Bloom (1970):

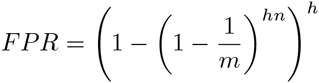

In the context of de Bruijn graph assembly, Bloom filter false positives have the effect of adding *k*-mers to the graph that are not present in the input sequencing reads. To address this issue, we have implemented a look-ahead mechanism to remove such *k*-mers from the graph. However, in order to confirm that Bloom filter false positives do not cause assembly artifacts, and to better understand the relationship between Bloom filter false positive rate, RAM usage, and running time, we conducted the following experiment.

Using the *C. elegans* dataset DRR008444, we assessed the effect of Bloom filter false positive rate on the NG50, number of misassemblies, run time, and peak memory usage of ABySS 2.0.0 (Fig. 1). As we increased the false positive rate from 1.9% to 20.7%, the NG50 remained roughly the same at 9600 bp, decreasing slightly as FPR reached 20% (Fig. 1A). Similarly, the number of misassemblies (9) remained constant across FPR values (Fig. 1B). As FPR was increased, the run time of ABySS 2.0.0 increased rapidly (Fig. 1C) while the peak memory usage decreased rapidly (Fig. 1D). These plots demonstrate that for this dataset we can trade off between memory usage and run time, with a FPR in the range of 5% - 12% giving both good memory and time performance. It also indicates that any FPR below 20% has no adverse effects on assembly quality in terms of contiguity or correctness.

**Figure 1:**
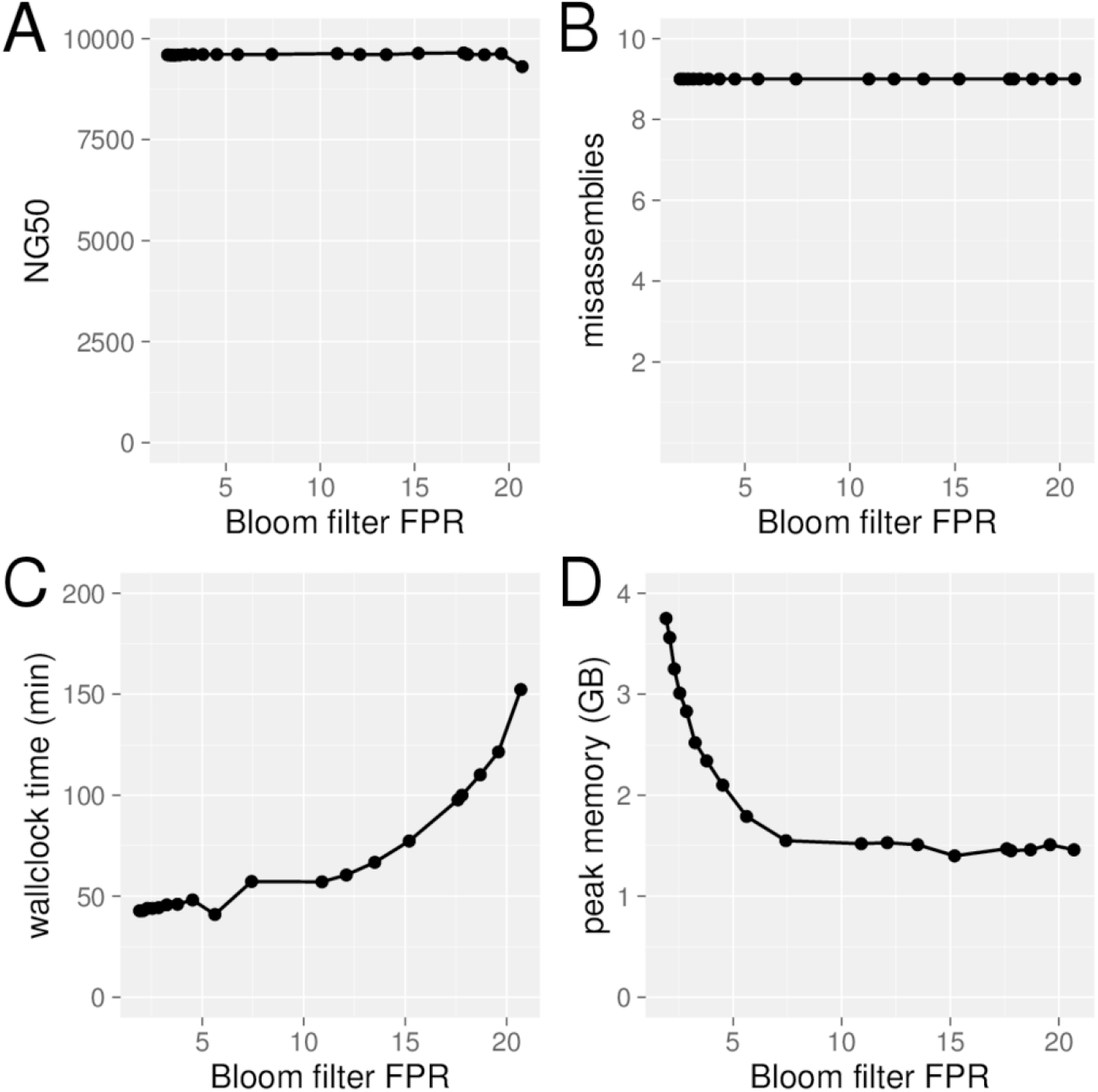
Effect of Bloom filter false positive rate (FPR) on ABySS 2.0.0 assemblies of the *C. elegans* DRR008444 dataset. (A) The assembly contiguity (NG50) is stable up to an FPR of 20%. (B) The number of misassemblies reported by QUAST (9) is stable with respect to FPR. (C) The assembly wallclock time increases with FPR and rises quickly when FPR exceeds 12%. (D) Peak memory usage drops quickly as FPR increases and levels out as FPR reaches 7.5%. From these results we conclude that a Bloom filter FPR in the range of 5-12% provides a good balance between assembly time and memory usage, without any detrimental effect on assembly quality.

### Assembler Comparison

To assess the performance of ABySS 2.0.0, we compared it with other leading assemblers for large genomes: ABySS 1.9.0 (Simpson et al. 2009), BCALM2 (Chikhi et al. 2016), DISCOVAR *de novo* 52488 (Weisenfeld et al. 2014), Minia 3.0 beta (Chikhi and Rizk 2013), SGA 0.10.14 (Simpson and Durbin 2011) and SOAPdenovo 2.04 (Luo et al. 2012). We conducted our comparison using a recent, publicly available human short read data set provided by the Genome in a Bottle (Zook et al. 2016) project. The Genome in a Bottle HG004 data was chosen for its deep 70X coverage of current (paired-end 250 bp) Illumina short read data and the availability of sequences from other platforms, including a 175X physical coverage jumping library (mate-pair) dataset (after trimming), 10X genomics Chromium data, and BioNano optical mapping data. Each of the assemblers included in the comparison was chosen due to its significant contributions towards the goal of scalable *de novo* assembly. ABySS facilitates large genome assemblies by distributing the de Bruijn graph across cluster nodes, and was the first software to assemble a human genome from short reads. The BCALM2 assembler introduces a novel method for partitioning the de Bruijn graph using minimizer hashing, which enables subsets of the graph to be assembled iteratively or in parallel. DISCOVAR *de novo* is a recent de Bruijn graph assembler for large genomes. Minia is the first assembler to employ a Bloom filter representation of the de Bruijn graph and uses a novel algorithm for eliminating Bloom filter false positives. SGA demonstrates the use of an FM-index (Simpson and Durbin 2011) as the core data structure for assembly, enabling detection of variable-length overlaps between reads with a low memory footprint. In addition to the aforementioned assemblers, we also attempted to include ALLPATHS-LG 52488 (Gnerre et al. 2010) and MaSuRCA 3.1.3 (Zimin et al. 2013) in our comparison. However, these assemblers did not run to completion on the target data set. ALLPATHS-LG 52488 (Gnerre et al. 2010) ran for one month and did not complete in that time. MaSuRCA 3.1.3 (Zimin et al. 2013) ran for five days and failed with a segmentation fault in the program `gatekeeper`.

We first compared the resource-efficiency of the assemblers by measuring their peak RAM and wallclock time (Fig. 2D, Table 4). Memory usage and run time varied wildly from 5 GB to 1.8 TB and 9 hours to 8 days. As expected given the succinct representation of the de Bruijn graph using Bloom filters, both Minia and ABySS 2.0.0 had memory footprints that were an order of magnitude smaller than other assemblers, with the exception of BCALM2, which both achieved the smallest memory footprint, by virtue of its novel partitioning strategy to constructing the de Bruijn graph, and completed the assembly in 9 hours, 8 hours of which was spent counting *k*-mers with DSK (Rizk et al. 2013). In contrast, DISCOVAR *de novo*, which achieves the best sequence contiguity, required 1.8 TB of memory and 8 days to complete.

**Figure 2:**
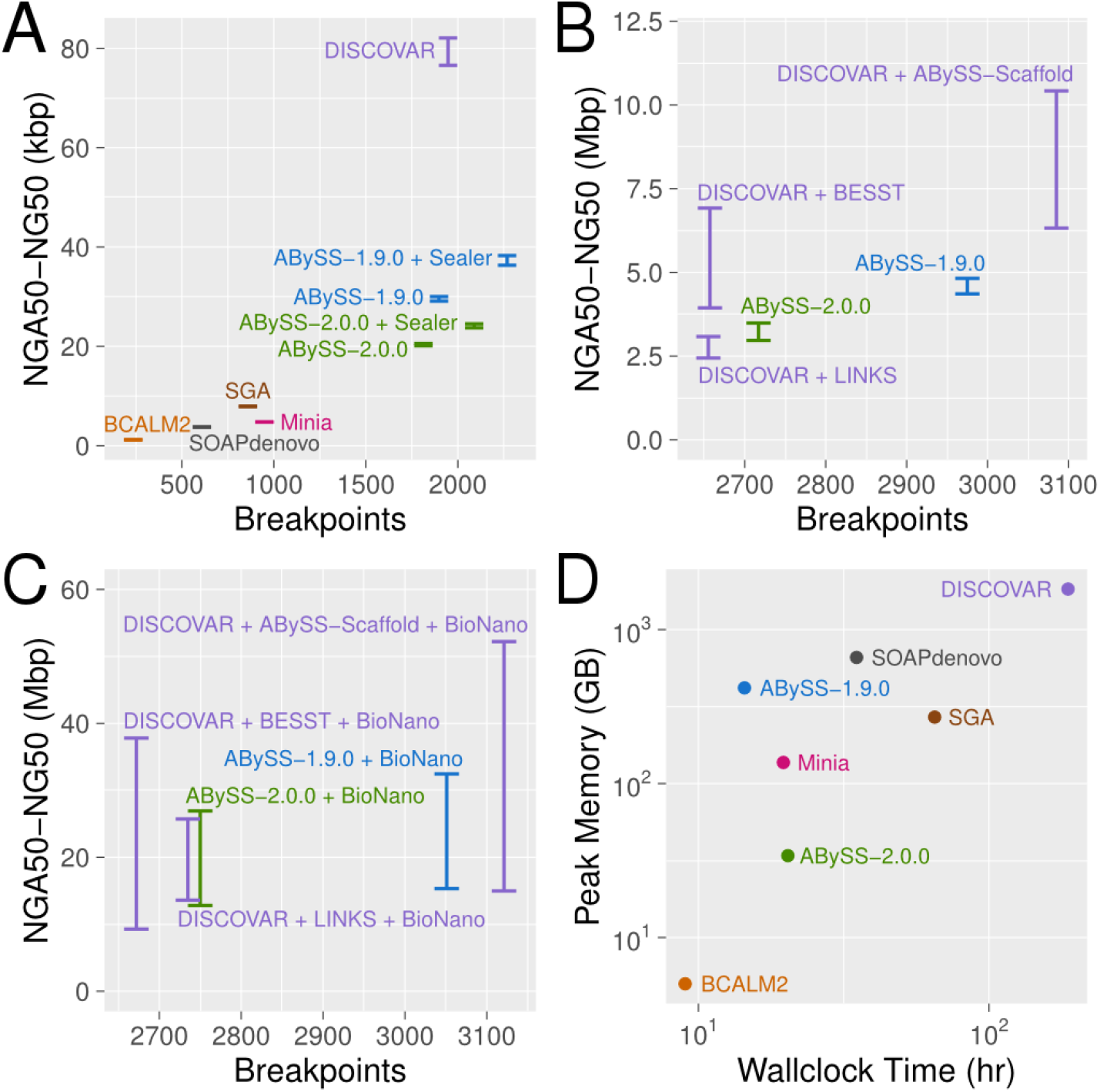
*De novo* assembly results for Genome in a Bottle HG004 human genome short read data with ABySS 1.9.0, ABySS 2.0.0, BCALM2, DISCOVAR, Minia, SOAPdenovo, and SGA. For panels A-C, on the Y axes we show the range of NGA50 to NG50 to indicate uncertainty caused by real genomic variants between individual HG004 and the reference genome (GRCh38). On the X axes, we show the number of breakpoints that occurred when aligning the sequences to the reference genome. Breakpoints are an indicator for the number of miassemblies but are also subject to uncertainty due to genomic variation between HG004 and the reference genome. (A) Contiguity and correctness metrics for contig sequences. For assemblies with scaffolding stages, the contigs were extracted by splitting the sequences at ‘N’ characters. (B) Contiguity and correctness metrics after scaffolding with mate pair (MPET) reads. The SOAPdenovo result for this plot was excluded as an outlier with an NGA50 (NG50) value of 103 kbp (172 kbp) and 10,610 breakpoints (C) Contiguity and correctness metrics after further scaffolding with BioNano optical mapping data, using BioNano’s IrysSolve software. (D) Peak memory usage and wallclock time for the assemblers. These wallclock times do not include the BioNano scaffolding stage, which was approximately 2 hours and did not affect peak RAM usage. The DISCOVAR wallclock time does not include the additional time for scaffolding with LINKS / BESST / ABySS-Scaffold.

**Table 4:** The peak memory usage and wall clock run time with 64 threads of the assemblies of GIAB HG004.

| Assembly | Memory (GB) | Time (h) |
| --- | --- | --- |
| ABYSS 1.9.0 | 418 | 14 |
| ABYSS 2.0.0 | 34 | 20 |
| DISCOVARdenovo 52488 | 1,832 | 187 |
| BCALM 2.0.0 | 5 | 9 |
| Minia 3.0-beta | 137 | 19 |
| SGA 0.10.14 | 270 | 65 |
| SOAPdenovo 2.04 | 659 | 35 |

We next compared the assemblies in terms of their contiguity and correctness (Fig. 2A-C, Tables 1-3). For contiguity assessment we calculated both NG50 and NGA50 using a genome size of 3,088,269,832 bp, whereas for correctness we aligned the contigs to the primary chromosome sequences of the human reference GRCh38 using BWA MEM 0.7.13 and counted the number of resulting break-points using `abyss-samtobreak -l500 -G3088269832`. As some assemblers such as BCALM2 and Minia only implement the first (de Bruijn graph) stage of assembly, we included comparisons for both the contig (Fig. 2A) and scaffold (Fig. 2B) stages, as applicable. To extract contig sequences from scaffolded assemblies, we split the sequences at occurrences of one or more ‘N’ characters. From the contig comparison, we observe that DISCOVAR achieves the highest sequence contiguity by a factor of approximately two (DISCOVAR NG50 of 82 kbp vs. ABySS + Sealer NG50 of 38 kbp). However, we also note that this result comes at the expense of an order of magnitude more time (8 days) and memory (1.8 TB) than the other assemblers, as shown in Fig. 2D. The NG50 of the ABySS 1.9.0 (30 kbp) and ABySS 2.0.0 (21 kbp) contigs noticeably exceeds those of BCALM2 (1 kbp) and Minia (5 kbp), primarily due to the additional use of paired-end information in ABySS. Comparing the contig results of the two ABySS assemblies, we note that the ABySS 2.0.0 assembly has slightly lower contiguity than ABySS 1.9.0 (21 kbp vs. 30 kbp). Upon investigation, we have observed that the main cause of this difference is the handling of low coverage regions. Whereas ABySS 1.9.0 retains all *k*-mers in the de Bruijn graph along with their counts, ABySS 2.0.0 discards *k*-mers with counts below a user-specified threshold, as discussed in Methods.

**Table 1:** The scaffold contiguity and number of breakpoints when aligned to GRCh38 using BWA MEM 0.7.13 of the assemblies of GIAB HG004 with BioNano scaffolding.

| Assembly | NG50 (Mbp) | NGA50 (Mbp) | Breakpoints |
| --- | --- | --- | --- |
| ABYSS 1.9.0 + BioNano | 32.5 | 15.3 | 3,051 |
| ABYSS 2.0.0 + BioNano | 26.9 | 12.8 | 2,750 |
| DISCOVAR + ABYSS-Scaffold + Bionano | 52.2 | 15.0 | 3,121 |
| DISCOVAR + LINKS + BioNano | 25.7 | 13.6 | 2,735 |
| DISCOVAR + BESST + BioNano | 37.8 | 9.3 | 2,672 |

**Table 2:** The scaffold contiguity and number of breakpoints when aligned to GRCh38 using BWA MEM 0.7.13 of the assemblies of GIAB HG004.

| Assembly | NG50 (Mbp) | NGA50 (Mbp) | Breakpoints |
| --- | --- | --- | --- |
| ABYSS 1.9.0 | 4.82 | 4.36 | 2,975 |
| ABYSS 2.0.0 | 3.49 | 2.97 | 2,717 |
| DISCOVAR + ABYSS-Scaffold | 10.42 | 6.32 | 3,085 |
| DISCOVAR + LINKS | 3.08 | 2.44 | 2,655 |
| DISCOVAR + BESST | 6.92 | 3.94 | 2,657 |
| SOAPdenovo 2.04 | 0.17 | 0.10 | 11,219 |

**Table 3:** The sequence contiguity and number of breakpoints when aligned to GRCh38 using BWA MEM 0.7.13 of the assemblies of GIAB HG004.

| Assembly | NG50 (kbp) | NGA50 (kbp) | Breakpoints |
| --- | --- | --- | --- |
| ABYSS 1.9.0 | 30.0 | 29.1 | 1,898 |
| ABYSS 1.9.0 + Sealer | 38.0 | 36.3 | 2,268 |
| ABYSS 2.0.0 | 20.6 | 20.1 | 1,813 |
| ABYSS 2.0.0 + Sealer | 24.5 | 23.7 | 2,089 |
| DISCOVARdenovo 52488 | 82.1 | 76.6 | 1,947 |
| BCALM 2.0.0 | 1.2 | 1.2 | 236 |
| Minia 3.0-beta | 4.8 | 4.7 | 949 |
| SGA 0.10.14 | 7.9 | 7.8 | 859 |
| SOAPdenovo 2.04 | 3.8 | 3.7 | 609 |

To further improve the contiguity of the ABySS contigs, we closed gaps in the assembly scaffolds with Sealer (Paulino et al. 2015). Sealer fills gaps by searching for a connecting path between gap flanks in the de Bruijn graph, and uses multiple *k*-mer sizes to maximize the probability of successfully finding a path. For the ABySS 1.9.0 assembly, Sealer closed 33,380 of 148,795 (22.4%) of scaffold gaps and increased the contig NG50 from 30 kbp to 38 kbp. For the ABySS 2.0.0 assembly, Sealer closed 33,533 of 213,480 (15.7%) scaffolds and increased the contig NG50 from 21 kbp to 25 kbp.

Comparing the contiguity/correctness of the scaffolded assemblies (Fig. 2B), we observe that the results of ABySS and DISCOVAR begin to converge, as do the results between the two versions of ABySS compared. As DISCOVAR does not provide its own mate-pair scaffolding algorithm, we augmented its assembly with three third-party scaffolders, ABySS-Scaffold, LINKS (Warren et al. 2015) and BESST (Sahlin et al. 2016), to enable a more direct comparison with ABySS. We note that there are significant differences between NG50 and NGA50 metrics in the scaffold plot, particularly in the case of the DISCOVAR + BESST assembly (3.9 Mbp vs. 6.9 Mbp). The NG50 is calculated under the assumption that all sequences are correctly assembled, whereas the NGA50 metric penalizes breakpoints when aligning the sequences to the reference genome. While on one hand the NG50 is an overly optimistic metric, on the other hand the NGA50 is an overly pessimistic metric because certain breakpoints may be attributed to real structural variation between the sequenced individual and the reference genome. For this reason, we show contiguity of the assemblies as a range between NGA50 and NG50, with the true unknown value lying somewhere in between.

After scaffolding the assemblies with Illumina mate-pair data, we performed an additional round of scaffolding using the BioNano optical mapping data and BioNano’s hybrid scaffolding tool hybridScaffold.pl. BioNano generates an optical map of the genome by fluorescently tagging occurrences of a particular endonuclease motif within long DNA molecules, resulting in a barcode-like pattern for each molecule. To perform the scaffolding, BioNano generates an analogous set of barcode patterns in silico for the sequences of the input assembly, and then aligns the two sets of bar codes. Applying BioNano scaffolding to the mate-pair-scaffolded sequences improves the NG50 and NGA50 contiguity metrics by roughly a factor of five across all assemblies (Fig. 2C), with NG50 reaching 52 Mbp with DISCOVAR + ABySS-Scaffold + BioNano. The distance between the NG50 and NGA50 values grows much larger at this stage of scaffolding, which we surmise is caused by a greater likelihood of encountering real sequence variation between the sequenced individual and the reference genome.

Our breakpoints metric of relative correctness between assemblies may be confounded by smaller real structural variations, especially as the assemblies become more contiguous. To this end, we investigated large-scale misassemblies (>10MB) and found only 2 major events within our ABySS 2.0.0 + BioNano scaffolds. One of these large scale events between chromosome 1 and 16 was identified in every assembly (Supplementary Fig. S1-S6), which indicates that the event may be real structural variant. The other large scale event is between chromosome 6 and 8 and is interestingly also found in the DISCOVAR + BESST + BioNano assembly (despite having fewer breakpoints and using an independent methodology), hence the relative correctness of the ABySS2 + BioNano assembly is still on par with the other assemblies.

### Scaffolding with Chromium Data

As the final step of our ABySS 2.0.0 assembly, we used the 10X Genomics Chromium data available for individual HG004 to further scaffold the Bionano assembly. The Chromium sequencing platform augments existing short read technologies by labeling reads that originate from the same long DNA molecule with a shared barcode sequence, also referred to as a read index. This labeling is achieved during library preparation by isolating long DNA molecules into droplets alongside gel beads containing the barcoding oligos. The read indices added by the Chromium protocol provide additional long-range grouping information for the short reads, which can be leveraged for scaffolding and other bioinformatics applications, such as phasing sequence variants.

To scaffold our assembly with the Chromium data, we aligned the Chromium reads to the input BioNano scaffolds with BWA MEM 0.7.13 and recorded the indices of the reads that aligned to each scaffold. As we were only interested in the read indices that joined scaffolds, we reduced noise by masking the interior portions of the input BioNano scaffolds with ‘N’ characters, preserving only the first/last 30 kbp of sequence in each scaffold, prior to aligning the Chromium reads. Using the information obtained from the read alignments, we constructed a graph representation of the relationships between scaffolds, using nodes to represent scaffolds and edge weights to represent the number of shared read indices between scaffolds. Finally, we supplied this graph as input to the LINKS (Warren et al. 2015) scaffolding algorithm to identify high-confidence paths within the graph and to output the corresponding scaffolds.

The Chromium scaffolding increased the scaffold NG50 of our ABySS 2.0.0 assembly from 26.9 Mbp to 41.9 Mbp. At this scale of contiguity, the largest scaffolds represent significant fractions of chromosome arms. In Fig. 3, we show the positions on the chromosomes of the 90 scaffolds larger than 3.2 Mbp that compose 90% of the genome. We note that many chromosome arms are reconstructed by 1 to 4 large scaffolds.

**Figure 3:**
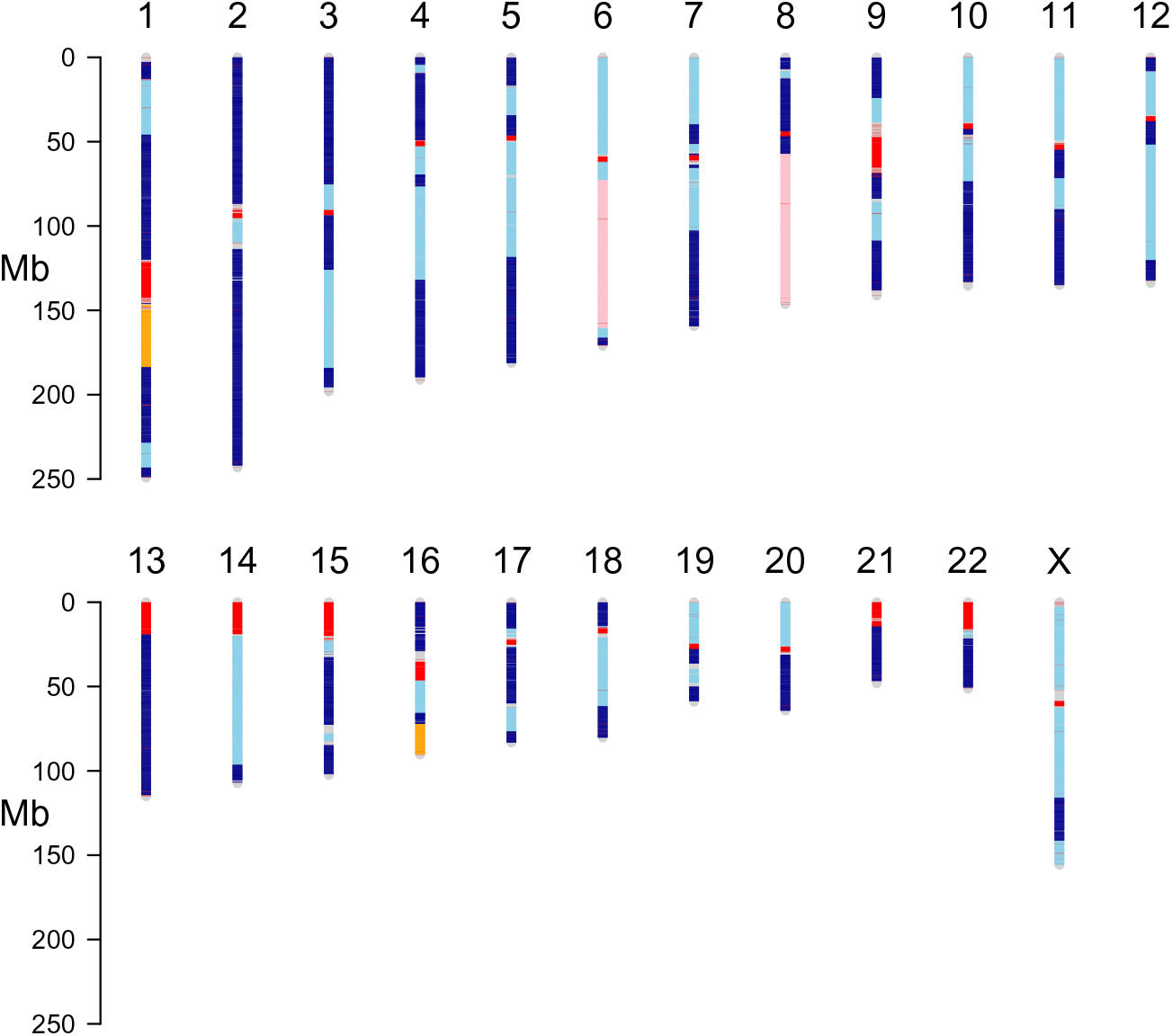
Contigs from the 90 scaffolds larger than 3.2 Mbp that compose 90% of the genome are aligned to GRCh38 using BWA-MEM 0.7.13. Contigs from the same scaffold are shown in the same shade of blue, and alternating shades of blue are used to distinguish between contigs from different scaffolds. Two translocations, t(1;16) and t(6;8), are shown in orange and pink. The segments of the genome that are not covered by alignments of the largest 90 scaffolds are shown in grey. Gaps in the reference genome, including centromeres and other heterochromatin, are shown in red.

**Figure 4:**
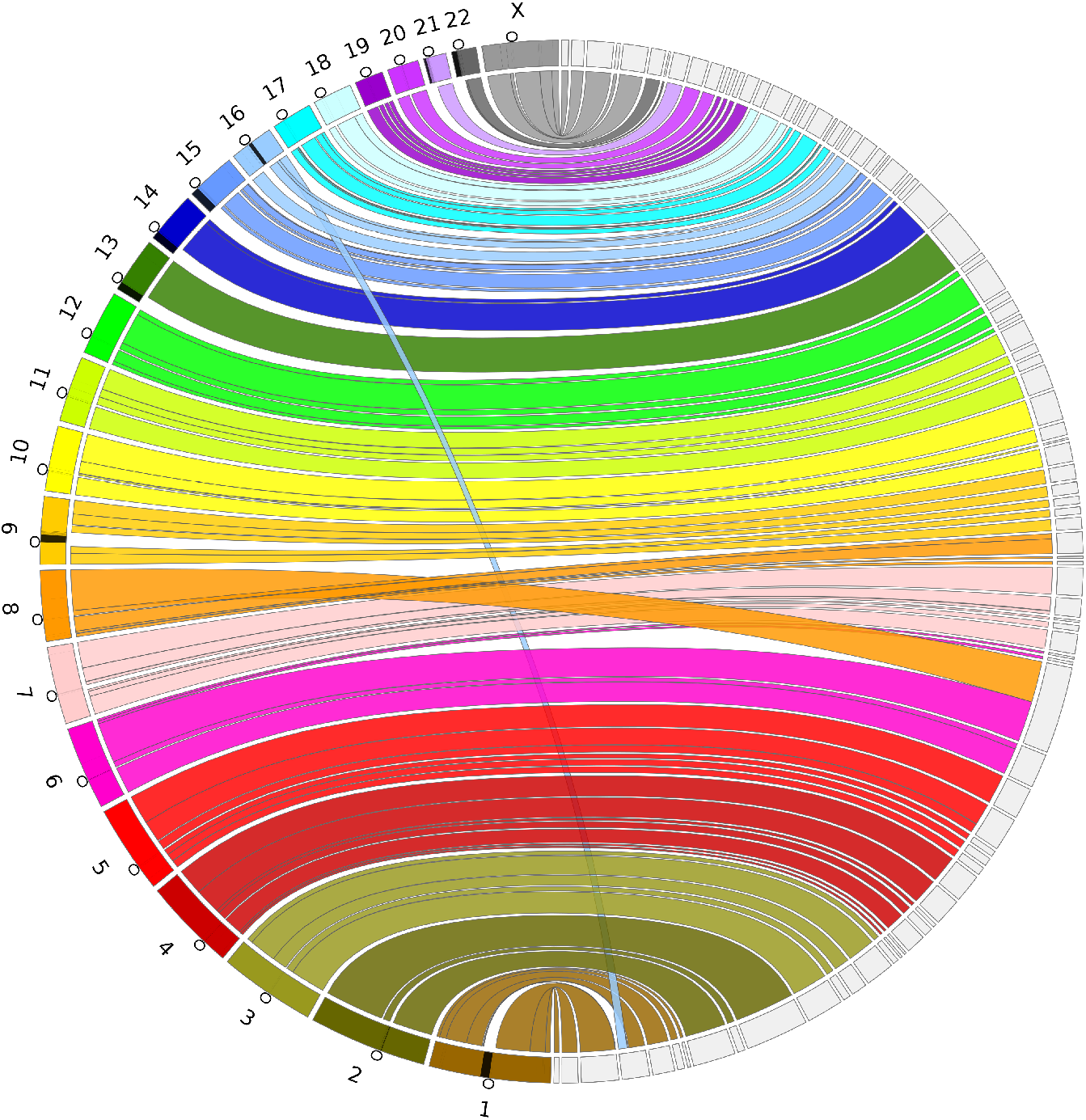
A Circos (Krzywinski et al. 2009) Assembly Consistency Plot. Scaftigs from the largest 89 scaffolds that compose 90% of the genome are aligned to GRCh38 using BWA-MEM 0.7.13. GRCh38 chromosomes are displayed on the left and the scaffolds on the right. Connections show the aligned regions between the genome and scaffolds. Contigs are included as a part of the same region if the are within 1Mbp of on either side of the connection, and regions shorter than 100Kbp are not shown. The black regions on the chromosomes indicate gaps in the reference and the circles indicate the centromere location on each chromosome.

## Discussion

The idiogram of Fig. 3 demonstrates that correct and highly-contiguous *de novo* assembly of human genomes is possible using current short read sequencing technologies combined with long range scaffolding techniques. While each of the scaffolding data types used here (mate-pair, BioNano, Chromium) are capable of increasing assembly contiguity by orders of magnitude on their own, our results demonstrate that these data are even more powerful when used in combination, also demonstrated by Mostovoy et al. (2016). In the human assembly we have described here, each scaffolding step feeds on the success of the previous assembly stages. Longer contig sequences improve the results of mate-pair scaffolding by allowing more mate-pairs to map to the contigs. Longer mate-pair scaffolds improve the BioNano scaffolding by allowing the optical map to align unambiguously to the mate-pair scaffolds; for this reason, BioNano recommends that the input assembly contains sequences of at least 100 kbp. Finally, longer BioNano scaffolds improve the Chromium scaffolding by resolving ambiguities in ordering and orientation of the scaffolds that are difficult to resolve using Chromium data alone.

Another observation that can be made from our assembler comparison is that, in spite of more than a decade of research and development related to de Bruijn graph assemblers, the memory and runtime efficiency of short read assemblers can still be greatly improved. This issue is particularly important for downstream studies that involve large numbers of *de novo* assemblies, such as human population studies, cancer genome studies, and clinical applications. The opportunity for improving the throughput of *de novo* assemblies is evident when comparing novel de Bruijn graph implementations such as Minia and BCALM2 against more mature assembly pipelines such as ABySS 1.9.0 and DISCOVAR (Fig. 2D). For example, the Minia assembler used only 137 GB RAM and required less than a day to run, whereas the equivalent DISCOVAR assembly used nearly 2 TB RAM and required more than a week to run. While Minia and BCALM2 did not match the results of ABySS and DISCOVAR in terms of assembly contiguity (Fig. 2A), we posit that this is due to the limited error removal of the implementations and not a fundamental limitation of the algorithms themselves. In the case of Minia, this hypothesis is borne out by the results of ABySS 2.0.0 (Fig. 2C), which employs a Bloom filter-based assembly approach similar to Minia, but achieves contiguity results that are on par with DISCOVAR and ABySS 1.9.0.

The assembly of long reads has yielded highly contiguous genome assemblies of human (Pendleton et al. 2015; Chin et al. 2016) and other organisms with sequence contiguity in the megabase range. Long read sequencing comes however at a cost premium. For applications that are cost sensitive, such as sequencing for diagnostic medicine, algorithms that exploit high-throughput short-read sequencing are valuable. We show that megabase scaffolds are achievable using short-read sequencing with one paired-end and one mate-pair library, and scaf-folds approaching the size of entire chromosome arms are possible with additional BioNano scaffolding. A remaining challenge is to improve the sequence contiguity of assemblies of short reads sequencing, which remain in the range of tens of kilobases, significantly shorter than the megabases achieved with the assembly of long read sequencing.

## Methods

### Bloom filter de Bruijn Graph Assembly

The first stage of the ABySS 2.0.0 assembly pipeline is a de Bruijn graph assembler that uses a compact, Bloom filter-based representation of the graph. The use of Bloom filters for *de novo* assembly was first demonstrated in Minia (Chikhi and Rizk 2013), and ABySS 2.0.0 builds on many aspects of that approach. The parts of our assembly algorithm that are novel with respect to Minia are: (i) the use of perfect reads to seed contig traversals, (ii) the use of look-ahead for error correction and elimination of Bloom filter false positives rather than a separate data structure, and (iii) the use of a new hashing algorithm, ntHash (Mohamadi et al. 2016), designed for processing DNA/RNA sequences efficiently. We describe these aspects of the algorithm in the course of our overall description below.

In the first step of the assembly, we load *k*-mers from the sequencing reads into a Bloom filter. These *k*-mers represent the set of nodes in a de Bruijn graph, even though we do not explicitly store the edges connecting the nodes. Instead, we discover edges at runtime by querying the Bloom filter for all four possible predecessors/successors of the current *k*-mer during the course of a graph traversal. Each possible successor (predecessor) corresponds to a single-base extension of the current *k*-mer to the right (left) by “A”, “C”, “G”, or “T”. To eliminate the majority of *k*-mers resulting from sequencing errors, we discard all *k*-mers with an occurrence count that is less than a user-specified threshold. We utilize a cascading Bloom filter to implement the filtering by *k*-mer count, as described in our previous work on Konnector (Vandervalk et al. 2014). Briefly, a cascading Bloom filter is a chained array of Bloom filters where each Bloom filter stores *k*-mers with a count that is one higher than the preceding Bloom filter. The procedure for inserting a *k*-mer into a cascading Bloom filter is to check for the presence of the *k*-mer in each Bloom filter in succession and to add the *k*-mer to the first Bloom filter where it is not already present. After all *k*-mers from the reads have been inserted, the last Bloom filter in the chain is then kept as the set of solid *k*-mers and the preceding Bloom filters are discarded.

In the second step of the assembly, we generate the unitig sequences by extending perfect reads left and right within the Bloom filter de Bruijn graph, where a read is considered to be perfect if it consists entirely of solid *k*-mers. The extension of the sequence continues left and right within the graph until either a dead end or a branching point is encountered. One complication of this approach is that Bloom filter false positives and recurrent sequencing errors will create branches in the graph that cause the sequence extension to end prematurely. To address this issue, we invoke an additional look-ahead step at each branching point, up to a distance of *k* nodes (Fig. 5). If the look-ahead step reveals that a branch is less than or equal to *k* nodes, it is considered to be a false branch and its existence is ignored. If, on the other hand, the branch point has two or more branches that are longer than *k* nodes then the extension is halted. The use of look-ahead incurs an additional computational cost to the graph traversal, but obviates the requirement for additional data structures to track false positives and error *k*-mers.

**Figure 5:**
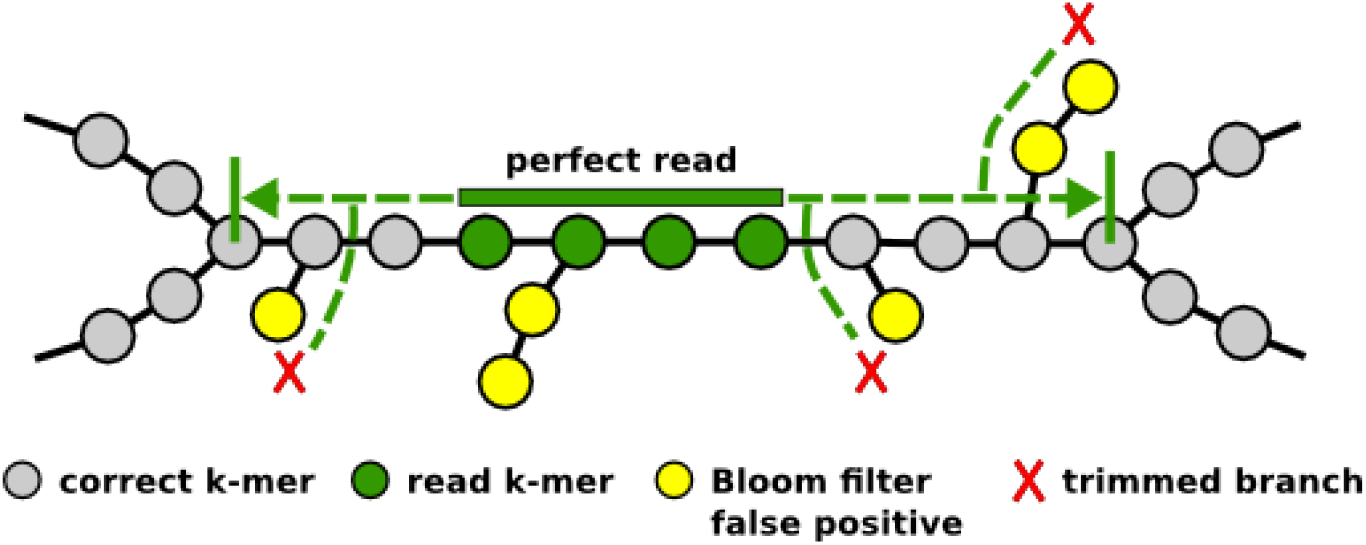
Extension of reads during Bloom filter de Bruijn graph assembly. A “perfect” read consisting only of solid *k*-mers is extended left and right within the de Bruijn graph until a branching point or dead-end is encountered. A look-ahead algorithm is employed to detect and ignore short branches due to Bloom filter false positives and/or recurrent read errors.

In the above steps, we use ntHash algorithm while working with the Bloom filter data structure. ntHash is an efficient hash method that computes the hash values for all consecutive *k*-mers in a DNA sequence recursively, in which the hash value for a *k*-mer is derived from the hash value of the previous *k*-mer. It is an adapted version of cyclic polynomial hashing to compute normal or canonical hash values for *k*-mers in DNA sequences efficiently. Further, ntHash provides a fast way to compute multiple hash values for a given *k*-mer, without repeating the whole hashing procedure for each value by few more operations. This is very useful for certain bioinformatics applications such as ABySS 2.0.0 that employs the Bloom filter data structure.

### Effect of Bloom Filter False Positive Rate

To assess the effects of the Bloom filter false positive rate on ABySS 2.0.0 assemblies, we ran multiple assemblies of the *C. elegans* N2 strain DRR008444 dataset (Illumina GA IIx sequencing of 2 × 100 bp reads of 300 bp fragments to 75 fold coverage) while varying the Bloom filter size from 250M to 3G with a step size of 250M. For example, the ABySS 2.0.0 assembly for a Bloom filter size of 250M was run with the command `abyss-pe c = 4 k = 64 H = 1 B = 250M in = 1 DRR008444_1.fastq DRR008444_2.fastq′`, where `c=4` specifies a minimum *k*-mer count threshold of 4, k = 64 specifies a *k*-mer size of 64, `H=1` specifies that the Bloom filter should use a single hash function. The runs for other Bloom filter sizes used the same parameter values with the exception of B (Bloom filter size).

For each assembly, we measured the wallclock time, peak memory usage, NG50, and number of misassemblies (Fig. 1). Wallclock time was measured with `/usr/bin/time`, while peak memory usage was determined by running the command `ps -eo pid,rss,vsz,cmd --width 100 --sort -vsz | awk ′NR==2′` in the background every 10 seconds to sample the top virtual-memory-consuming process. We used QUAST 3.2 (Gurevich et al. 2013) to calculate the NG50 length and number of major misassemblies, using the *C. elegans* Bristol N2 strain as the reference genome (NCBI BioProject PRJNA158). The false positive rates corresponding to each Bloom filter size were obtained from the ABySS log files. All assemblies were run with 12 threads on an isolated machine with 48GB RAM and two Xeon X5650 CPUs.

### Assembler Comparison

#### Experimental Sequencing Data

The Genome in a Bottle (GIAB) project (Zook et al. 2016) sequenced seven individuals using a large variety of sequencing technologies. We downloaded the Illumina WGS 2 × 250 bp paired-end sequencing data and the Illumina 6 kbp mate-pair sequencing data of the Ashkenazi mother (HG004).

We removed adapters from the mate-pair reads using NxTrim 0.4.0 (O’Connell et al. 2014) with parameters `--norc --joinreads --preserve-mp`, which also classifies the reads as mate-pair, paired-end, single-end or unknown. We discarded the reads classified as either paired-end or single-end, and used the reads classified as mate-pair and unknown for scaffolding, which are comprised primarily of mate-pair reads originating from large fragments.

We corrected sequencing errors in the reads using the tool BFC (Li 2015) with the parameter `-s3G`. We constructed the hash table of trusted *k*-mers using the paired-end reads and used this hash table to correct both the paired-end and mate-pair reads.

#### Human Assemblies

We assembled the GIAB HG004 data set using ABySS 1.9.0 (Simpson et al. 2009), ABySS 2.0.0, ALLPATHS-LG 52488 (Gnerre et al. 2010), BCALM 2.0.0 (Chikhi et al. 2016), DISCOVARdenovo 52488 (Weisenfeld et al. 2014), MaSuRCA

3.1.3 (Zimin et al. 2013), Minia 3.0 beta (Chikhi and Rizk 2013), SGA 0.10.14 (Simpson and Durbin 2011), SOAPdenovo 2.0.4 (Luo et al. 2012). We assembled with each tool the paired-end reads corrected by BFC. The mate-pair reads categorized by NxTrim and corrected by BFC and were used for scaffolding when applicable for that assembler. We scaffolded the DISCOVARdenovo assembly using both BESST 2.2.4 (Sahlin et al. 2016) and LINKS 1.6.1 (Warren et al. 2015).

Most software used in these analyses was installed from the Homebrew-Science software collection using Linuxbrew (Jackman and Birol 2016) with the command `brew install abyss allpaths-lg bcalm bfc bwa discovardenovo masurca nxtrim samtools seqtk sga soapdenovo`. The development version of ABySS-2.0.0 used in the comparison was compiled from the `bloom-abyss-preview` tag: https://github.com/bcgsc/abyss/tree/ bloom-abyss-preview. Minia 3.0 beta and LINKS 1.6.1 were installed manually, as the versions currently available in Linuxbrew are 2.0.3 and 1.5.1 respectively. The Python package `besst` was installed using `pip install besst`. The script used to assemble and analyze the data is a Makefile script available online at https://github.com/sjackman/giab/blob/1.0/Makefile.

We assembled the paired-end and mate-pair reads using ABySS 1.9.0 (Simpson et al. 2009) with the command `abyss-pe name=hsapiens np=64 k=144 q=15 v=-v l=40 s=1000 n=10 S=1000-10000 N=15 mp6k_de=--mean mp6k_n=1 lib=pe400 pe400=$(<pe400.in)` mp=mp6k mp6k=$(mp6k + unknown.in) where the files pe400.in and mp6k + unknown. in are lists of the locations of compressed FASTQ files.

We assembled the paired-end and mate-pair reads using ABySS 2.0.0 with the command abyss-pe name=hsapiens np=64 k=144 q=15 v=-v l=40 s=1000 n=10 S=1000-10000 N=15 B=26G H=4 c=3 mp6k_de=--mean mp6k_n=1 lib=pe400 pe400=$(<pe400.in) mp=mp6k mp6k=$(mp6k + unknown.in). The parameters for ABySS-2.0.0 were identical to those for ABySS-1.9.0, with the exception of the Bloom filter specific parameters `B=26G H=4 c=3`, which specify the total memory allocated to the Bloom filters, the number of Bloom filter hash functions, and the number of cascading Bloom filter levels, respectively. We determined the values for total memory size (B) and number of hash functions (H) by counting distinct 144-mers with KMC2 (Deorowicz et al. 2015) and targeting a false positive rate of 5% for the first level of the cascading Bloom filter. We deemed 5% to be a suitable upper bound for Bloom filter FPR based on the results of our *C. elegans* experiment above, which indicated good performance in the range of 5-12% FPR. We determined the optimal number of cascading Bloom filter levels by running assemblies with `c=2, c=3`, and `c=4`, and choosing the assembly with highest NG50 and lowest number of breakpoints.

We assembled the paired-end and mate-pair reads using ALLPATHS-LG 52488 (Gnerre et al. 2010) with the command `PrepareAllPathsInputs.pl DATA_DIR=${PWD} PLOIDY=2 HOSTS=32; RunAllPathsLG PRE=. REFERENCE_NAME=. DATA_SUBDIR=. RUN=allpaths SUBDIR=run` and the configuration files in_libs.csv and in_groups.csv shown in supplementary material and online at https://github.com/sjackman/giab/tree/1.0/allpaths-lg.

We assembled the paired-end reads using BCALM 2.0.0 (Chikhi et al. 2016) with the commands `bcalm -in pe400.in -out hsapiens-unitigs -k 63-abundance 5 -nb-cores 64; bglue -in hsapiens-unitigs.h5 -out hsapiens-unitigs -k 63`. The largest value of *k* supported by BCALM 2.0.0 is 63.

We assembled the paired-end reads using DISCOVARdenovo 52488 (Weisenfeld et al. 2014) with the command DiscovarDeNovo MAX_MEM_GB=2500 READS=@pe600.in OUT_DIR=./hsapiens and scaffolded this assembly using three standalone scaffolding tools: ABySS-Scaffold 1.9.0 with the command abyss-pe `name=hsapiens mp=mp6k j=64 k=200 l=40 s=500 S=500-5000 N=15 mp6k_de=--mean mp6k_n=1 mp6k=$(mp6k + unknown.in)` scaffolds, BESST 2.2.4 (Sahlin et al. 2016) with the command `runBESST --orientation rf -c hsapiens-scaffolds.fa -f mp6k.bam -o .`, and LINKS 1.6.1 (Warren et al. 2015) with the command ***todo*** @rwarren.

We assembled the paired-end and mate-pair reads using MaSuRCA 3.1.3 (Zimin et al. 2013) with the command `./masurca config.txt; ./assemble.sh` and the configuration file `config.txt` shown in supplementary material and online at https://github.com/sjackman/giab/blob/1.0/masurca/config.txt. The script `assemble.sh` is generated by `masurca` itself.

We assembled the paired-end reads using Minia 3.0 beta (Chikhi and Rizk 2013) with the command `minia -in pe400.in -abundance-min “auto”-kmer-size 128 -nb-cores 64`. The largest value of *k* supported by this pre-release version was 128.

We assembled the paired-end reads using SGA 0.10.14 (Simpson and Durbin 2011) with the commands sga preprocess --pe-mode=2 hsapiens.fa.gz > hsapiens.preprocess.fa; sga index -t 64 -a ropebwt hsapiens.preprocess.fa; sga filter -t 64 hsapiens.preprocess.fa; sga overlap -t 64 hsapiens.preprocess.filter.pass. -m 125; sga assemble -o hsapiens hsapiens.preprocess.filter.pass.asqg.gz.

We assembled the paired-end and mate-pair reads using SOAPdenovo 2.0.4 (Luo et al. 2012) with the command `SOAPdenovo-127mer all -K 95 -R –p 63 -s hsapiens.config` and the configuration file `hsapiens.config` shown in supplementary material and online at https://github.com/sjackman/giab/blob/ 1.0/soapdenovo/hsapiens.config.

We used the BioNano optical map to scaffold the ABySS 1.9.0, ABySS 2.0.0 and DISCOVARdenovo 52488 assemblies, scaffolded with ABySS-Scaffold, BESST and LINKS, using IrysSolve 2.1.5063 with the command line `hybridScaffold.pl -n hsapiens-scaffolds.fa -b EXP_REFINEFINAL1_q.cmap -c hybridScaffold_config_aggressive.xml -B2 -N2 -o bionano -x -y -m all.bnx -q optArguments_human.xml -e AJmother_autoNoise1.err` according to the document “Theory Of Operation: Hybrid Scaffolding” available online at http://bionanogenomics.com/wp-content/ uploads/2016/04/30073-Rev-A-Hybrid-Scaffolding-Theory-of-Operations.pdf. The configuration files are used unmodified as distributed by BioNano Genomics and available online at https://github.com/sjackman/giab/tree/1.0/bionano.

We used 10x Genomics Chromium data to scaffold the ABySS 2.0.0 + BioNano scaffolds with ARCS (unpublished) and LINKS (Warren et al. 2015). The version of ARCS used in the paper is available from: https:

//github.com/bcgsc/arcs/tree/arcs-prerelease. First we aligned the Chromium reads to the ABySS 2.0.0 + BioNano scaffolds using bwa mem with default settings. Next we ran ARCS with the command `arcs -f hsapiens-scaffolds.fa -a human-alignments.fof -s 98 -g 50000 -c 5 -l 5 -m 50-1000 -d 0 -e 30000 -i 16 -v 1` where `hsapiens-scaffolds.fa` contained the sequences to further scaffold and `human-alignments.fof` was a file of alignment file filenames. Lastly we ran the commands `python makeTSVfile.py hsapiens-scaffolds.fa.scaff_s98_c5_original.gv human_c5.tigpair_checkpoint.tsv hsapiens-scaffolds.fa; LINKS -f hsapiens-scaffolds.fa -s empty.fof -b human_c5 -l 5 -a 0.3` to order and orient the scaffolds. The script `makeTSVfile.py` can be found at https://github.com/sarahyeo/giab.

## Data Access

The accession number of the Ashkenazi mother is NIST HG004 NA24143 SRS823307.

The Illumina WGS 2 × 250 bp paired-end sequencing data may be downloaded from https://github.com/genome-in-a-bottle/giab_data_indexes/blob/master/ AshkenazimTrio/sequence.index.AJtrio_Illumina_2 × 250bps_06012016. The 35 experiment accession numbers are SRX1726894 through SRX1726928, and the 35 sequencing run accession numbers are SRR3440461 through SRR3440495.

The Illumina 6 kbp mate-pair sequencing data may be downloaded from https://github.com/genome-in-a-bottle/giab_data_indexes/blob/master/ AshkenazimTrio/sequence.index.AJtrio_Illumina_6kb_matepair_wgs_ 08032015. The two experiment accession numbers are SRX1388736 and SRX1388737, and the two sequencing run accession numbers are SRR2832452 and SRR2832453.

The BioNano optical map `EXP_REFINEFINAL1_q.cmap` may be downloaded from https://github.com/genome-in-a-bottle/giab_data_indexes/blob/master/

AshkenazimTrio/alignment.index.AJtrio_BioNano_xmap_cmap_GRC37_ 10012015.

The 10x Genomics Chromium data may be downloaded from https://github. com/genome-in-a-bottle/giab_data_indexes/blob/master/AshkenazimTrio/ alignment.index.AJtrio_10Xgenomics_ChromiumGenome_GRCh37_ GRCh38_06202016.

## Acknowledgements

The research presented here was funded by the National Human Genome Research Institute of the National Institutes of Health (under award number R01HG007182), with additional support provided by Intel, Genome Canada, Genome British Columbia, and the British Columbia Cancer Foundation. The content is solely the responsibility of the authors and does not necessarily represent the official views of the National Institutes of Health or other funding organizations. We would also like to thank Martin Krzywinski for his help with the data visualization in the idiogram and Circos figures.

## References

Bloom BH. 1970. Space/time trade-offs in hash coding with allowable errors. Communications of the ACM 13: 422–426.

Chaisson MJP, Huddleston J, Dennis MY, Sudmant PH, Malig M, Hormozdiari F, Antonacci F, Surti U, Sandstrom R, Boitano M, et al. 2014. Resolving the complexity of the human genome using single-molecule sequencing. Nature 517: 608–611. http://dx.doi.org/10.1038/nature13907.

Chikhi R, Limasset A, Jackman S, Simpson JT, Medvedev P. 2014. On the representation of de bruijn graphs. Research in Computational Molecular Biology 35–55. http://dx.doi.org/10.1007/978-3-319-05269-4_4.

Chikhi R, Limasset A, Medvedev P. 2016. Compacting de bruijn graphs from sequencing data quickly and in low memory. Bioinformatics 32: i201–i208. http://dx.doi.org/10.1093/bioinformatics/btw279.

Chikhi R, Rizk G. 2013. Space-efficient and exact de bruijn graph representation based on a bloom filter. Algorithms for Molecular Biology 8: 1.

Chin C-S, Peluso P, Sedlazeck FJ, Nattestad M, Concepcion GT, Clum A, Dunn C, O’Malley R, Figueroa-Balderas R, Morales-Cruz A, et al. 2016. Phased diploid genome assembly with single molecule real-time sequencing. http://dx.doi.org/10.1101/056887.

Deorowicz S, Kokot M, Grabowski S, Debudaj-Grabysz A. 2015. KMC 2: Fast and resource-frugal k-mer counting. Bioinformatics 31: 1569–1576.

Gnerre S, MacCallum I, Przybylski D, Ribeiro FJ, Burton JN, Walker BJ, Sharpe T, Hall G, Shea TP, Sykes S, et al. 2010. High-quality draft assemblies of mammalian genomes from massively parallel sequence data. Proceedings of the National Academy of Sciences 108: 1513–1518. http://dx.doi.org/10.1073/pnas.1017351108.

Gurevich A, Saveliev V, Vyahhi N, Tesler G. 2013. QUAST: Quality assessment tool for genome assemblies. Bioinformatics 29: 1072–1075. http://dx.doi.org/10.1093/bioinformatics/btt086.

Jackman SD, Birol I. 2016. Linuxbrew and homebrew for cross-platform package management [v1; not peer reviewed]. F1000 Research 5(ISCB Comm J): 1795 (poster). http://dx.doi.org/10.7490/f1000research.1112681.1.

Krzywinski M, Schein J, Birol I, Connors J, Gascoyne R, Horsman D, Jones SJ, Marra MA. 2009. Circos: An information aesthetic for comparative genomics. Genome research 19: 1639–1645.

Lander ES, Linton LM, Birren B, Nusbaum C, Zody MC, Baldwin J, Devon K, Dewar K, Doyle M, FitzHugh W, et al. 2001. Initial sequencing and analysis of the human genome. Nature 409: 860–921. http://dx.doi.org/10.1038/35057062.

Ley T, Miller C, Ding L, Raphael B, Mungall A, Robertson A, Hoadley K, Triche TJ, Laird P, Baty J, et al. 2013. Genomic and epigenomic landscapes of adult de novo acute myeloid leukemia. N Engl J Med 368: 2059–2074. http://dx.doi.org/10.1056/NEJMoa1301689.

Li H. 2015. BFC: Correcting illumina sequencing errors. Bioinformatics 32: 2885–2887. http://dx.doi.org/10.1093/bioinformatics/btv290.

Luo R, Liu B, Xie Y, Li Z, Huang W, Yuan J, He G, Chen Y, Pan Q, Liu Y, et al. 2012. SOAPdenovo2: An empirically improved memory-efficient short-read de novo assembler. GigaSci 1. http://dx.doi.org/10.1186/2047-217X-1-18.

Mohamadi H, Chu J, Vandervalk BP, Birol I. 2016. NtHash: Recursive nucleotide hashing. Bioinformatics. http://dx.doi.org/10.1093/bioinformatics/btw397.

Morin RD, Mungall K, Pleasance E, Mungall AJ, Goya R, Huff RD, Scott DW, Ding J, Roth A, Chiu R, et al. 2013. Mutational and structural analysis of diffuse large b-cell lymphoma using whole-genome sequencing. Blood 122: 1256–1265. http://dx.doi.org/10.1182/blood-2013-02-483727.

Mose LE, Wilkerson MD, Hayes DN, Perou CM, Parker JS. 2014. ABRA: Improved coding indel detection via assembly-based realignment. Bioinformatics 30: 2813–2815. http://dx.doi.org/10.1093/bioinformatics/btu376.

Mostovoy Y, Levy-Sakin M, Lam J, Lam ET, Hastie AR, Marks P, Lee J, Chu C, Lin C, Džakula Ž, et al. 2016. A hybrid approach for de novo human genome sequence assembly and phasing. Nat Meth 13: 587–590. http://dx.doi.org/10.1038/nmeth.3865.

Nagarajan N, Pop M. 2013. Sequence assembly demystified. Nature Reviews Genetics 14: 157–167. http://dx.doi.org/10.1038/nrg3367.

O’Connell J, Schulz-Trieglaff O, Carlson E, Hims MM, Gormley NA, Cox AJ. 2014. NxTrim: Optimized trimming of illumina mate pair reads. http://dx.doi.org/10.1101/007666.

Paulino D, Warren RL, Vandervalk BP, Raymond A, Jackman SD, Birol I. 2015. Sealer: A scalable gap-closing application for finishing draft genomes. BMC bioinformatics 16: 230.

Pendleton M, Sebra R, Pang AWC, Ummat A, Franzen O, Rausch T, Stütz AM, Stedman W, Anantharaman T, Hastie A, et al. 2015. Assembly and diploid architecture of an individual human genome via single-molecule technologies. Nat Meth 12: 780–786. http://dx.doi.org/10.1038/nmeth.3454.

Pevzner PA, Tang H, Waterman MS. 2001. An eulerian path approach to dna fragment assembly. Proceedings of the National Academy of Sciences 98: 9748–9753.

Pugh TJ, Morozova O, Attiyeh EF, Asgharzadeh S, Wei JS, Auclair D, Carter SL, Cibulskis K, Hanna M, Kiezun A, et al. 2013. The genetic landscape of high-risk neuroblastoma. Nat Genet 45: 279–284. http://dx.doi.org/10.1038/ng.2529.

Rizk G, Lavenier D, Chikhi R. 2013. DSK: K-mer counting with very low memory usage. Bioinformatics 29: 652–653. http://dx.doi.org/10.1093/bioinformatics/btt020.

Roberts KG, Morin RD, Zhang J, Hirst M, Zhao Y, Su X, Chen S-C, Payne-Turner D, Churchman ML, Harvey RC, et al. 2012. Genetic alterations activating kinase and cytokine receptor signaling in high-risk acute lymphoblastic leukemia. Cancer Cell 22: 153–166. http://dx.doi.org/10.1016/j.ccr.2012.06.005.

Sahlin K, Chikhi R, Arvestad L. 2016. Assembly scaffolding with pe-contaminated mate-pair libraries. Bioinformatics 32: 1925–1932. http://dx.doi.org/10.1093/bioinformatics/btw064.

Simpson JT, Durbin R. 2011. Efficient de novo assembly of large genomes using compressed data structures. Genome Research 22: 549–556. http://dx.doi.org/10.1101/gr.126953.111.

Simpson JT, Wong K, Jackman SD, Schein JE, Jones SJ, Birol I. 2009. ABySS: A parallel assembler for short read sequence data. Genome Research 19: 1117–1123. http://dx.doi.org/10.1101/gr.089532.108.

Vandervalk BP, Jackman SD, Raymond A, Mohamadi H, Yang C, Attali DA, Chu J, Warren RL, Birol I. 2014. Konnector: Connecting paired-end reads using a bloom filter de bruijn graph. Bioinformatics and Biomedicine (BIBM), 2014 IEEE International Conference on 51–58.

Warren RL, Yang C, Vandervalk BP, Behsaz B, Lagman A, Jones SJM, Birol I. 2015. LINKS: Scalable, alignment-free scaffolding of draft genomes with long reads. GigaSci 4. http://dx.doi.org/10.1186/s13742-015-0076-3.

Weisenfeld NI, Yin S, Sharpe T, Lau B, Hegarty R, Holmes L, Sogoloff B, Tabbaa D, Williams L, Russ C, et al. 2014. Comprehensive variation discovery in single human genomes. Nat Genet 46: 1350–1355. http://dx.doi.org/10.1038/ng.3121.

Yip S, Butterfield YS, Morozova O, Chittaranjan S, Blough MD, An J, Birol I, Chesnelong C, Chiu R, Chuah E, et al. 2011. Concurrent cic mutations, idh mutations, and 1p/19q loss distinguish oligodendrogliomas from other cancers. The Journal of Pathology 226: 7–16. http://dx.doi.org/10.1002/path.2995.

Zimin AV, Marcais G, Puiu D, Roberts M, Salzberg SL, Yorke JA. 2013. The masurca genome assembler. Bioinformatics 29: 2669–2677. http://dx.doi.org/10.1093/bioinformatics/btt476.

Zook JM, Catoe D, McDaniel J, Vang L, Spies N, Sidow A, Weng Z, Liu Y, Mason CE, Alexander N, et al. 2016. Extensive sequencing of seven human genomes to characterize benchmark reference materials. Sci Data 3: 160025. http://dx.doi.org/10.1038/sdata.2016.25.

